# Estimating fruit tree growth curves in breeding field using fragmented longitudinal data: An application to citrus hybrid seedlings

**DOI:** 10.1101/2025.01.19.632888

**Authors:** Soh Kimura, Mai F. Minamikawa, Keisuke Nonaka, Tokurou Shimizu, Hiroyoshi Iwata

## Abstract

Vegetative and reproductive growth in fruit trees is interconnected, and analyzing this relationship can provide valuable insights into fruit quality. However, characterizing vegetative growth through growth models is challenging because of the difficulty in obtaining longitudinal data, given the slow growth rate. In breeding fields, in contrast, seedlings of different ages are planted, allowing for simultaneous measurements that yield a dataset resembling longitudinal data with missing values --termed “fragmented longitudinal data.” Because longitudinal data are obtained from a single measurement, they can potentially shorten the period required for growth curve estimation. Bayesian nonlinear models offer advantages in estimating curves from incomplete data. In this study, we generated fragmented longitudinal data using genome data with 45,929 markers from 624 citrus hybrid seedlings and applied a Bayesian nonlinear model to explore its potential. We also incorporated genomic information into the model to assess the impact of the estimation accuracy. Our simulations indicated that the Bayesian nonlinear model’s ability to interpolate missing values significantly improved the estimation performance. At best, the mean square error of the parameter characterizing the later growth stage was reduced by 84.3 mm^2^. Although the improvement from incorporating genomic information was modest, it still surpassed models that lacked genomic data. We also predicted the curves of untested individuals using the estimated parameters. Although the prediction accuracy of each parameter measured by the correlation coefficient was lower than 0.5, one parameter consistently showed a better accuracy. Further research is required to reveal the advantages of integrating genomic data for better predictions.

## Introduction

Vegetative growth in fruit trees is closely linked to reproductive growth, acting as both a source and sink. Understanding this relationship has been a key focus for researchers because it directly influences fruit quality^1^. While physiological research has traditionally dominated the study of this relationship^2,3^, quantitative analysis of vegetative traits, such as trunk diameter and crown width, along with subsequent correlation studies, has highlighted the significant role of vegetative growth in determining fruit quality^4–7^. Additionally, these traits have been shown to correlate with the juvenile period and inbreeding depression^8–10^. With the advent of next-generation sequencing, these analyses are expected to expand to the genetic level. However, the complex growth patterns of trees and the dynamic nature of their phenotypes over time make the genetic analysis of these traits highly time-dependent. Therefore, it is essential to evaluate the evolving growth patterns and align genetic analyses accordingly.

The coexistence of trees of different ages in the same field presents specific challenges for genetic analyses, particularly for traits that are strongly influenced by time. In fruit tree breeding, orchards often contain a mix of candidates selected at various growth stages. For instance, in Japanese Citrus breeding, approximately 1,000 seedlings obtained from various crosses are grafted onto trifoliate orange rootstocks annually and assessed for over 20 traits, including fruit quality and disease resistance^11,12^. Given that selection periods are typically under 8 years and new seedlings are planted annually, breeding orchards contain trees of diverse ages from various crosses. To analyze genetic variations in trees of various ages simultaneously, it is necessary to link data from different growth stages using a growth model and evaluate their growth patterns based on the parameters of that model.

Growth models such as the logistic and Gomperz models are effective in describing tree growth trajectories using a minimal number of time-independent parameters^13^. Their nonlinearity allows them to outperform linear models, such as polynomial regression, in terms of requiring fewer interpretable parameters and providing more stable extrapolation^14^, which has led to extensive studies evaluating their applicability^15^. Growth models have been used to better understand growth patterns of fruits^16^. The parameters estimated from these models, which serve as time-independent indices of growth, are often used as target traits for genomic selection^17,18^ or genome-wide association study (GWAS)^19,20^. Prior research has highlighted both the challenge of analysis based on time-dependent indices and the potential of the growth model, proposing QTL analysis based on stage-dependent indices derived from these models^21^. Unlike cross-sectional data, which can be obtained from a single measurement, longitudinal data, which are necessary for estimating growth curves, require repeated sampling, placing a significant burden on researchers. This burden is particularly high for trees, which have slow growth rates and require measurements over several years. Recently, unmanned aerial vehicles (UAVs), such as drones, have been used to measure phenotypic data and show promise in reducing this burden^22,23^. However, despite the anticipated reduction in measurement load owing to advancements in UAV technology, acquiring longitudinal data will continue to be challenging owing to the time required.

The use of an age-mixed field along with nonlinear mixed-effect models (NLMEMs) and Bayesian nonlinear models offers a potential solution for this challenge. In an age-mixed field, trees of different ages are planted together, allowing for simultaneous measurement of all individuals. This results in a dataset similar to longitudinal data, which we refer to as “fragmented longitudinal data.” Although these data contained substantial missing values for each tree (genotype), they were collected over a relatively short time span. If accurate growth curves can be estimated from these fragmented longitudinal data, the overall measurement time can be significantly reduced.

The nonlinear mixed effect model (NLMEM), which combines the characteristics of a nonlinear model and a mixed-effect model, is frequently employed in the literature to handle incomplete longitudinal data^24,25^. NLMEM includes both fixed and random effects, where fixed parameters are shared among all subjects and random parameters are unique to each individual^26^. Because individual-specific values of random parameters are estimated based on a covariance structure derived from all individuals in the dataset, each individual tree (genotype) can “borrow” information from the others, allowing for the imputation of missing values^13^. Bayesian nonlinear mixed models extend NLMEMs by assuming that all model parameters, not only random effects, follow specific distributions^27^, which further aid in imputing missing values^28^. Therefore, we expect that NLMEM and Bayesian nonlinear modeling will play crucial roles in estimating growth curves from fragmented longitudinal data. Recent studies have incorporated genomic information into these models^29,30^. As growth parameters are genetically controlled to some extent, incorporating genomic information into these models may improve parameter estimation accuracy. Moreover, when genomic information is considered, the growth curves of untested individuals can be predicted based on genomic data in the context of genomic prediction and selection^31^.

In this study, we applied Bayesian nonlinear models to estimate growth curve parameters using fragmented longitudinal data collected from age-mixed fields without the need for time-consuming, labor-intensive repeated measurements. In addition, we evaluated whether incorporating genomic information into the models could improve the accuracy of parameter estimation. To validate the potential of the model and the role of genomic information, we used real genomic data from citrus breeding materials and simulated their vegetative growth curves. In these simulations, we compare the estimation accuracy of several methods and scenarios using fragmented longitudinal data. Furthermore, we assessed genomic prediction accuracy for untested individuals. Thus, we identified the potential of these methods to estimate growth curves based on short-term data for materials with varying age structures.

## Results

### Generating fragmented longitudinal data from genome data

The main purpose of this study was to estimate and predict growth curves from fragmented longitudinal data obtained from an age-mixed field. To accurately evaluate the estimation and prediction performance, rather than using raw data, fragmented longitudinal data were generated from artificially created longitudinal data based on the following measurement design: In this experiment, we assumed that measurements were conducted once a year over a 2-year period in a citrus breeding field. During the first measurement, trees grafted 2, 4, and 6 years ago, which were categorized as the youngest, middle, and oldest cohorts, respectively, were measured. The same trees were measured again in the second year, resulting in 6 years of longitudinal data with 2 data points per individual (Fig. 1A).

**Fig 1.**
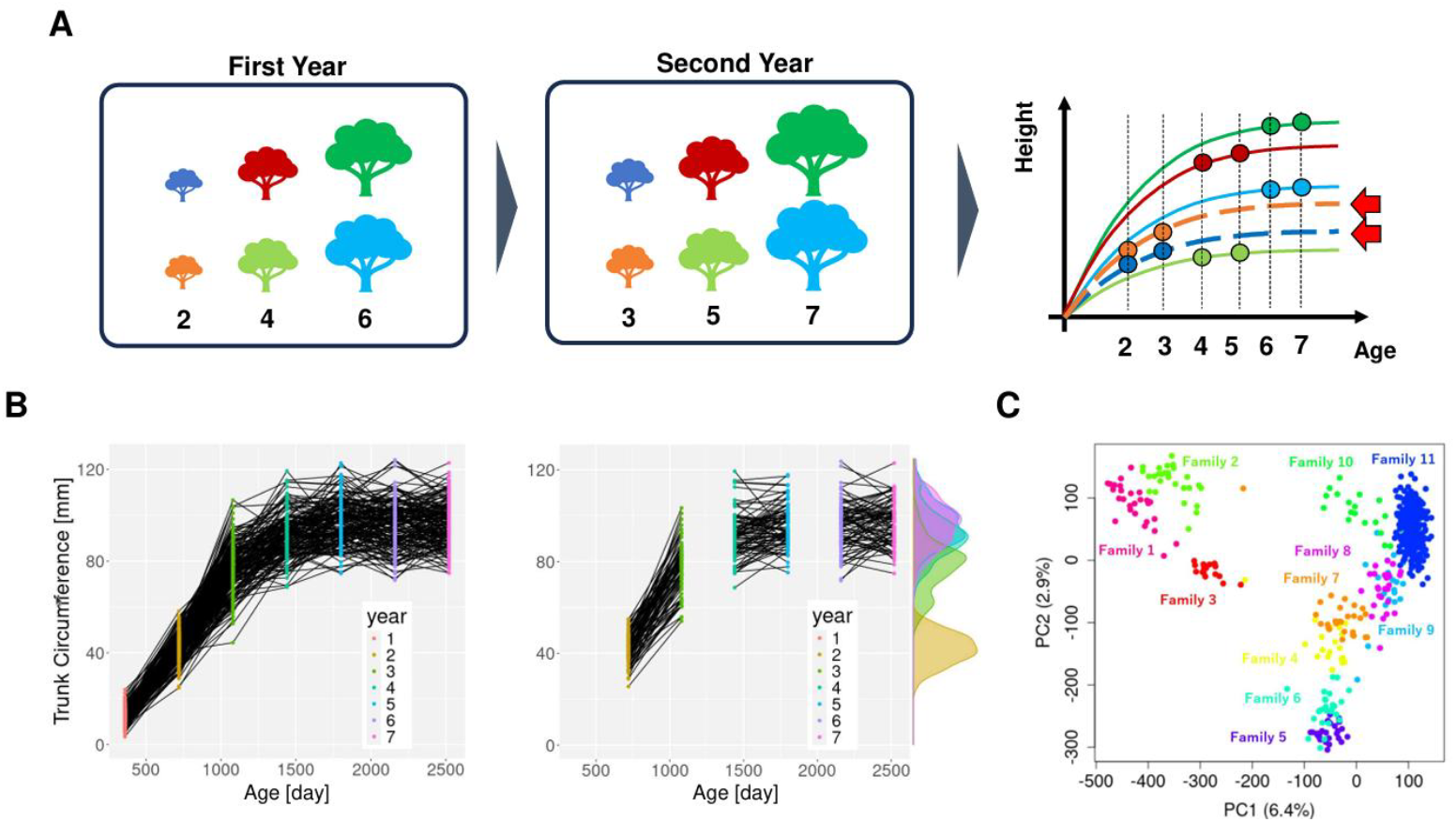
Measurement design and prepared fragmented longitudinal data. (A) Detailed depiction of fragmented longitudinal data sampling and experimental scenario. The individuals with the same color are identical in the two left images, and the numbers below the trees represent the years after grafting. Measurements were conducted over 2 years for the same individuals, resulting in fragmented longitudinal data from years 2 to 7 within 2-year period. (B) Generated longitudinal data and the corresponding fragmented longitudinal data for multiple-family groups. The dots represent sampling data and are colored based on the year of sampling. (C) Principal component analysis of genomic data: all individuals were distinguished by 11 colors, corresponding to their respective families. The numbers on each axis shown in parentheses indicate the contribution rates.

To investigate the influence of population structure on the estimation and prediction performance, two different longitudinal datasets were generated based on population composition: one from a single-family population and the other from a population of 11 families (Table 1). We refer to the former as the “single-family group” and the latter as the “multiple-family group.” During the generation of fragmented longitudinal data, the multiple-family group was further divided into two different groups; “multiple-family group I” and “multiple-family group II” depending on the allocation pattern of individuals. In single-family and multiple-family Group I, all individuals were randomly assigned to three cohorts. However, in actual breeding situations, the same cross is not repeated over multiple years, resulting in all individuals derived from the same cross concentrated within a specific year. To evaluate the estimation and prediction performance under these conditions, individuals in multiple-family group II were allocated to each cohort by family. Family allocations were determined randomly for each simulation trial (iteration).

**Table 1.** Number of individuals in each group.

| Group | Family number |  |  |  |  |  |  |  |  |  |  |
| --- | --- | --- | --- | --- | --- | --- | --- | --- | --- | --- | --- |
|  | 1 | 2 | 3 | 4 | 5 | 6 | 7 | 8 | 9 | 10 | 11 |
| single-family group | 0 | 0 | 0 | 0 | 0 | 0 | 0 | 0 | 0 | 0 | 266 |
| multiple-family group | 27 | 28 | 18 | 19 | 27 | 27 | 27 | 27 | 20 | 19 | 27 |

The generated and prepared fragmented longitudinal data are shown in Fig. 1B. Despite the dynamic fluctuation caused by the measurement noise in each curve, the overall patterns of both the generated and fragmented longitudinal data follow a typical logistic growth curve. The phenotype data for each year formed a distribution that resembled a normal distribution owing to both genomic and residual variance. Although the position of the distribution shifted upward over time, the distributions of the middle and oldest cohorts overlapped significantly, indicating that the generated longitudinal data began to converge at an early stage.

The genetic structures of all individuals used to generate the longitudinal data were visualized using principal component analysis (Fig. 1C), with 11 families represented by 11 distinct colors. Family 11, which formed a single-family group, was concentrated in the top-right section of the graph, whereas the 11 families that comprised the multiple-family group were widely dispersed across the graph, reflecting their greater genetic diversity.

### Estimation accuracy of future/past growth using a nonlinear model

To assess the difference in the estimation performance depending on the section of the estimated growth curve, two different scenarios were devised (Fig. 2A). In “Scenario 1,” the growth curve of the youngest cohort was estimated, while “Scenario 2” focuses on the oldest cohort, allowing the evaluation estimation performance for future. (younger), and past (older) growth. In Scenario 1, although the fragmented longitudinal data for the youngest cohort lacked information on later growth, the middle and oldest cohorts contained longitudinal data covering the later stages of growth. If the growth curve estimation for the youngest cohort can leverage information from the later growth of other cohorts, estimation performance is expected to improve. Similarly, estimating younger age growth using data from the oldest cohort is challenging. However, the estimation performance improves if the growth curve of the oldest cohort is jointly estimated with those of the other two cohorts. To validate this assumption, three estimation methods were compared for each scenario: Method 1, in which the growth curve of the youngest/oldest cohort was estimated from only the youngest/oldest cohort data without genomic information; Method 2, in which the growth curve of the youngest/oldest cohort was estimated using data from all cohorts without genomic information; and Method 3, in which the growth curve of the youngest/oldest cohort was estimated using data from all cohorts with genomic information, which was expected to yield the highest estimation accuracy.

**Fig 2.**
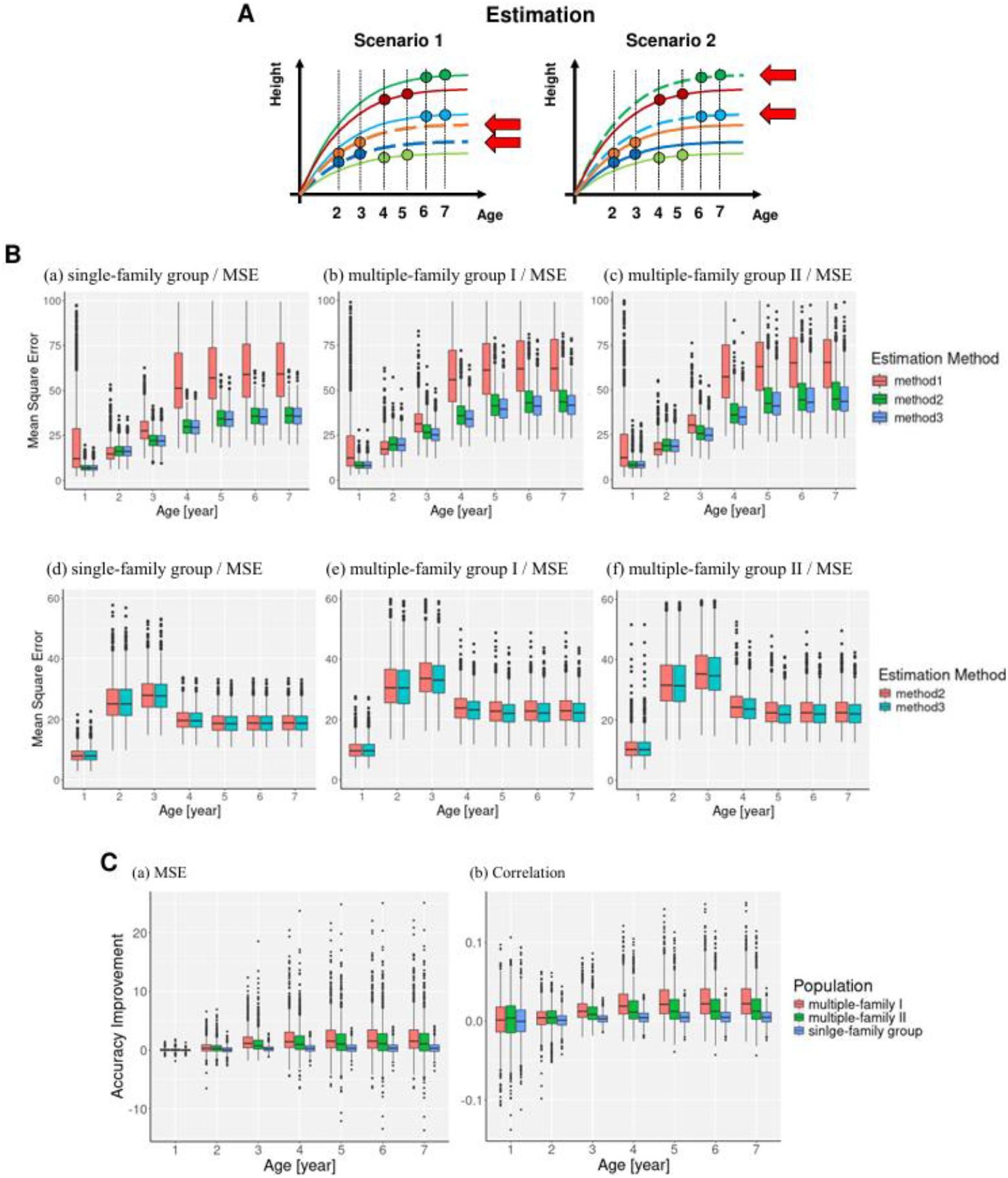
Estimation scenario and estimation performance. (A) In scenario 1, the later growth of the youngest cohort was estimated using three different methods, and their estimation performance was compared. Similarly, in scenario 2, the old cohort was targeted, and the estimation performance for younger growth was compared (B) The upper three images (a), (b), and (c) show the results of scenario 1, whereas the lower three images (d), (e), and (f) represent the results of scenario 2. Within each scenario, the left image corresponds to the single-family group, middle to the multiple-family group I, and right to the multiple-family group II. In each image, the estimation error of the growth curve was evaluated annually using the three different methods, each represented by distinct colors. Boxplots are not shown when growth curve could not be predicted. (C) Time variation of accuracy improvement: The x-axis represents the age of the growth curve, and the y-axis represents the accuracy improvement at each age. Estimation performance was evaluated using mean square error in the left image (a) and correlation coefficient in the right image (b). Accuracy improvement was compared across groups, with different colors representing each group.

For the estimation, the fragmented longitudinal data were fitted to a logistic model with three parameters: *A, B*, and *C*. Although parameter *A* characterizes the later growth, parameters *B* and *C* determine the dynamics of the initial growth stage. The similarity between the estimated and true curves was evaluated at ages 1–7 years for each scenario using both the mean square error (MSE) and Pearson’s correlation coefficient (correlation coefficient), as shown in Fig. 2B.

In scenario 1, where the later growth of the youngest cohort was estimated, the MSEs of methods 2 and 3 decreased substantially compared with method 1, particularly at ages 4–7 years across all groups. While the medians of the MSE of method 3 at ages 4 to 7 years were consistently smaller than those of method 2, the degree of superiority varied depending on the group (Fig. 2B (a), (b), and (c)). Similarly, the estimation performance evaluated by the correlation coefficient at ages 4–7 years showed the worst results for Method 1 and the best for Method 3 (Supplementary Fig. 1 (a), (b), and (c)). It should be noted that the MSE for Method 1 reached as high as 8000 mm^2^, but values beyond 100 mm^2^ were not shown on the y-axis owing to its scaling.

In scenario 2, where the younger growth of the oldest cohort was estimated, the function used to calculate the initial value for the Bayesian nonlinear model function did not work properly; therefore, no growth curve was obtained for method 1. The MSE of both methods were high, particularly at ages two and three (Fig. 2B (d), (e), and (f)), and the correlation coefficient was relatively low at ages one and two (Supplementary Fig. 1 (d), (e), and (f)), indicating imputation limitations. Similar to Scenario 1, the median MSE in Method 3 was consistently smaller than that in Method 2, and the median and interquartile range of the MSE in the single-family group were better than those in the other two groups.

Although the advantage of incorporating genomic information to improve estimation performance at ages 4–7 years was confirmed to be significant in scenario 1, the magnitude of this advantage was influenced by the population structure. To further investigate the relationship between population and the magnitude of improvement, we defined an index called “accuracy improvement.” This index was calculated for each simulation iteration as follows: (estimation accuracy in Method 3) minus (estimation accuracy in Method 2) for the correlation coefficient, and (estimation error in Method 2) minus (estimation error in Method 3) for the MSE. Because the estimation performance for Methods 2 and 3 was calculated simultaneously using the same created longitudinal data in each simulation iteration, taking the difference in each iteration makes it possible to evaluate the improvement in estimation performance by considering the genomic information more clearly. As a result, the accuracy improvement was found to be positive in most iterations and showed clear differences among groups in both the MSE and correlation coefficient. Multiple-family group I showed the best results, whereas the single-family group performed the worst (Fig. 2C).

The estimation performance of the latent parameters *A, B*, and *C*, which control the overall growth behavior, was also investigated, focusing on multiple-family group II in scenario 1 to identify the cause of accuracy improvement. Because the results of Method 1 sometimes included outliers that skewed the mean value, the estimation performance was summarized using the median of 1000 iterations instead of the mean, as shown in Table 2. In scenario 1, the MSEs of all parameters were confirmed to improve in methods 2 and 3, with a notable improvement in methods *A* and *B* when genomic information was considered. However, in Scenario 2, although the MSE of parameters *A* and *B* were higher in Method 3 than in Method 2, the magnitude of the difference was not as pronounced as that in Scenario 1.

**Table 2.**
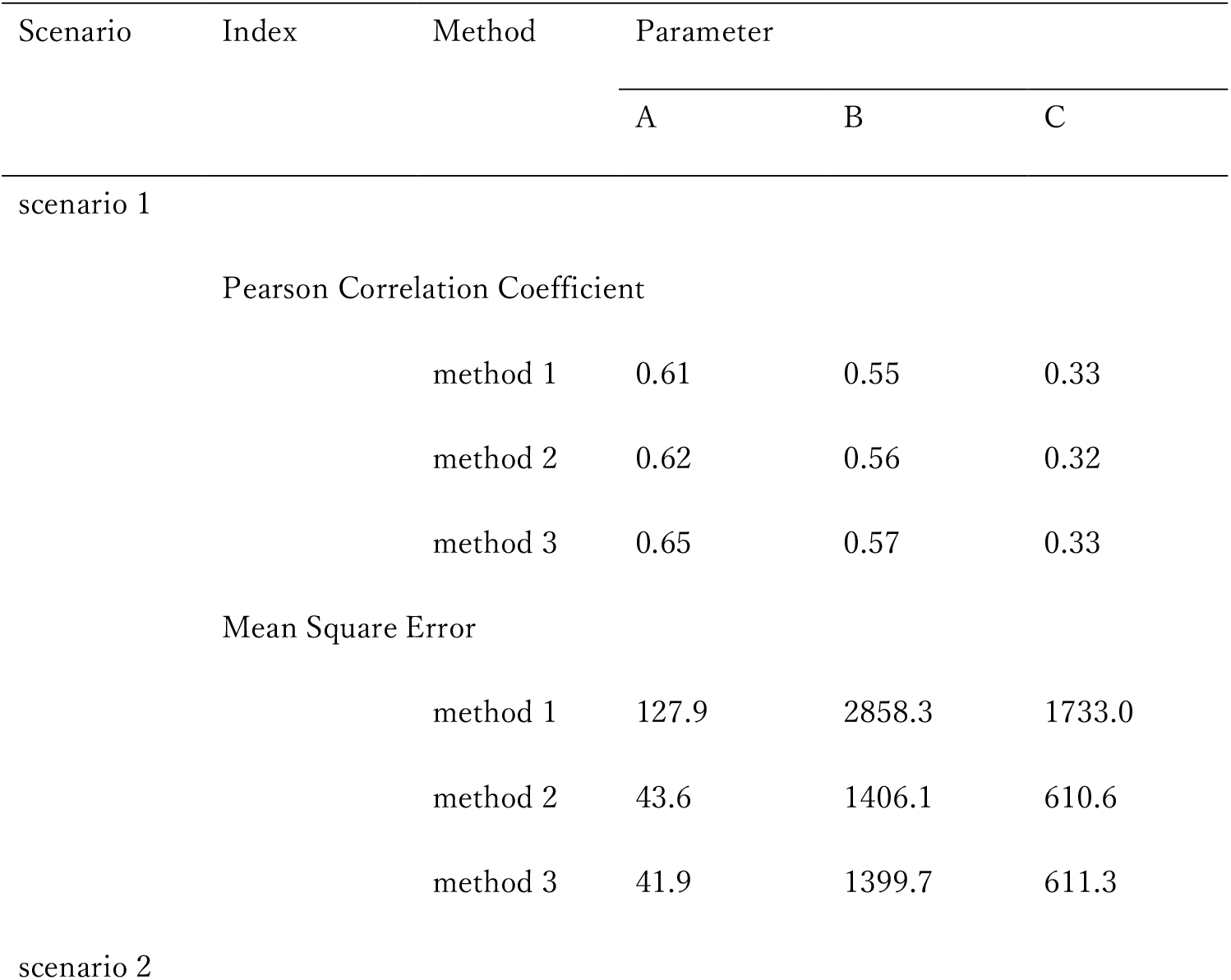
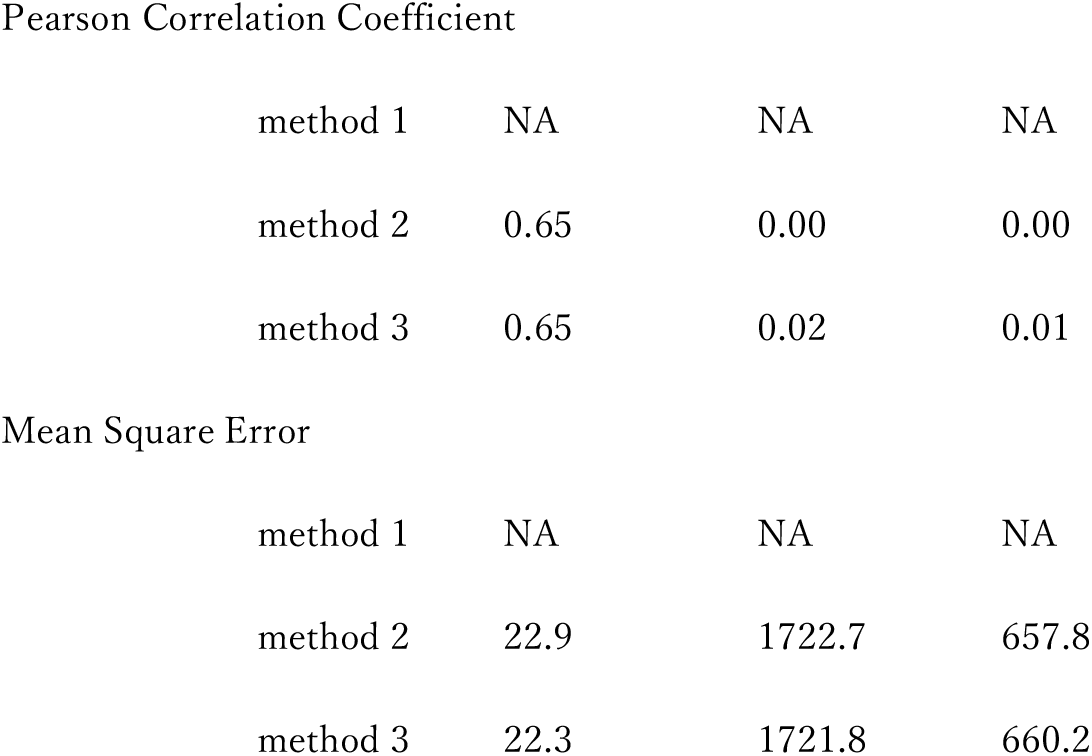
Estimation performance of each parameter.

| Scenario | Index | Method | Parameter |  |  |
| --- | --- | --- | --- | --- | --- |
|  |  |  | A | B | C |
| scenario 1 |  |  |  |  |  |
|  | Pearson Correlation Coefficient |  |  |  |  |
|  |  | method 1 | 0.61 | 0.55 | 0.33 |
|  |  | method 2 | 0.62 | 0.56 | 0.32 |
|  |  | method 3 | 0.65 | 0.57 | 0.33 |
|  | Mean Square Error |  |  |  |  |
|  |  | method 1 | 127.9 | 2858.3 | 1733.0 |
|  |  | method 2 | 43.6 | 1406.1 | 610.6 |
|  |  | method 3 | 41.9 | 1399.7 | 611.3 |
| scenario 2 |  |  |  |  |  |

Pearson Correlation Coefficient
|  |  |  |  |
| --- | --- | --- | --- |
| method 1 | NA | NA | NA |
| method 2 | 0.65 | 0.00 | 0.00 |
| method 3 | 0.65 | 0.02 | 0.01 |

Mean Square Error
|  |  |  |  |
| --- | --- | --- | --- |
| method 1 | NA | NA | NA |
| method 2 | 22.9 | 1722.7 | 657.8 |
| method 3 | 22.3 | 1721.8 | 660.2 |

### Predicting untested progeny using the model trained on incomplete longitudinal data

To date, estimation of the growth curve has been limited to individuals with phenotypic data. However, by incorporating the genomic information, it was possible to predict the growth curves of untested individuals (Fig. 3A). To assess the prediction performance of the model trained on fragmented longitudinal data, the prediction performances of latent parameters *A, B*, and *C* were evaluated using cross-validation. The prediction accuracy (correlation coefficient) of parameter *A* was higher than those of parameters B and C across all groups, with multiple-family group II showing the lowest accuracy for all parameters (Fig. 3B).

**Fig 3.**
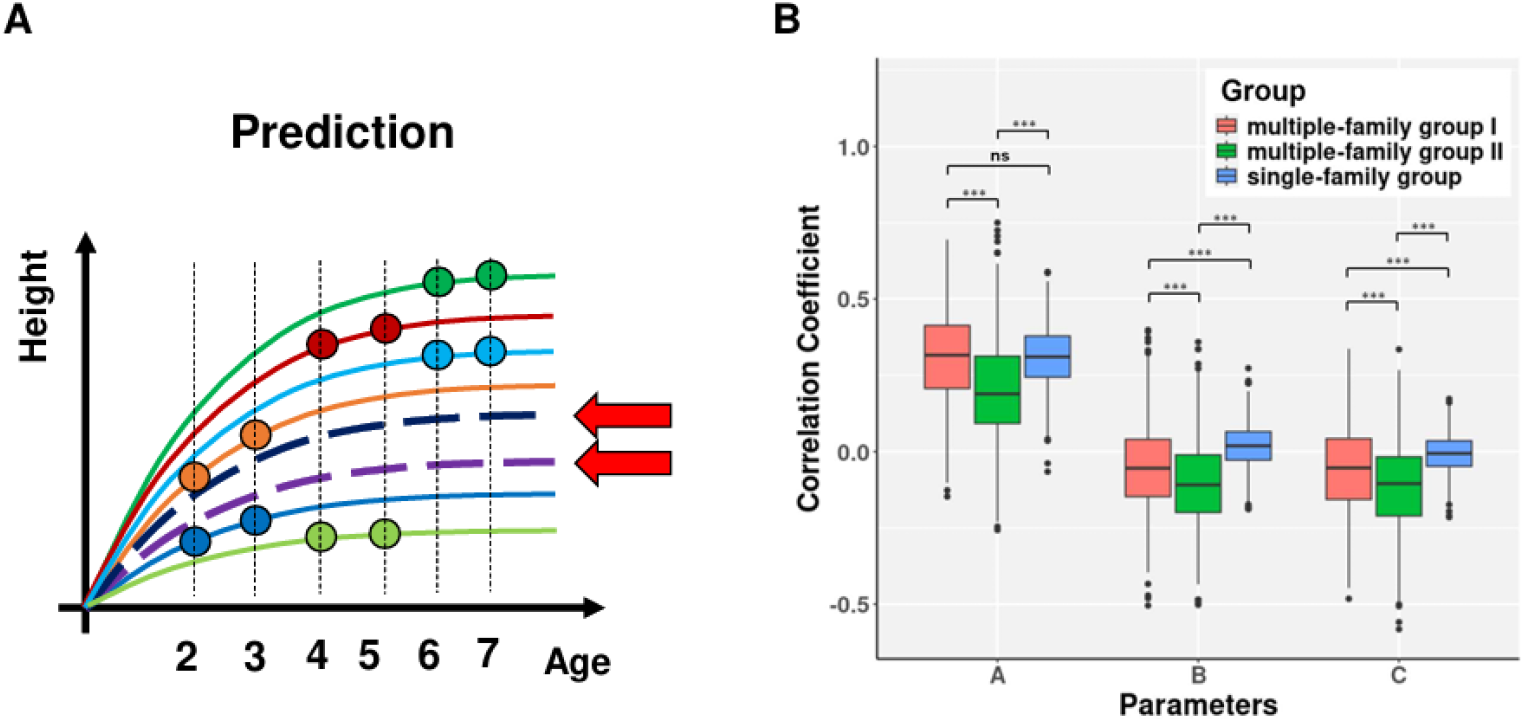
Prediction scenario and prediction performance. (A) Individuals without phenotype were predicted using the model developed with fragmented longitudinal data. (B) Prediction accuracy of all parameters in the logistic model. For each parameter, the results for three different groups are represented by distinct colors. Statistical significance is indicated by asterisk: *** for *p* ≤ *0*.*00*1 and ns for *p* > 0.05.

**Supplementary Fig 1.**
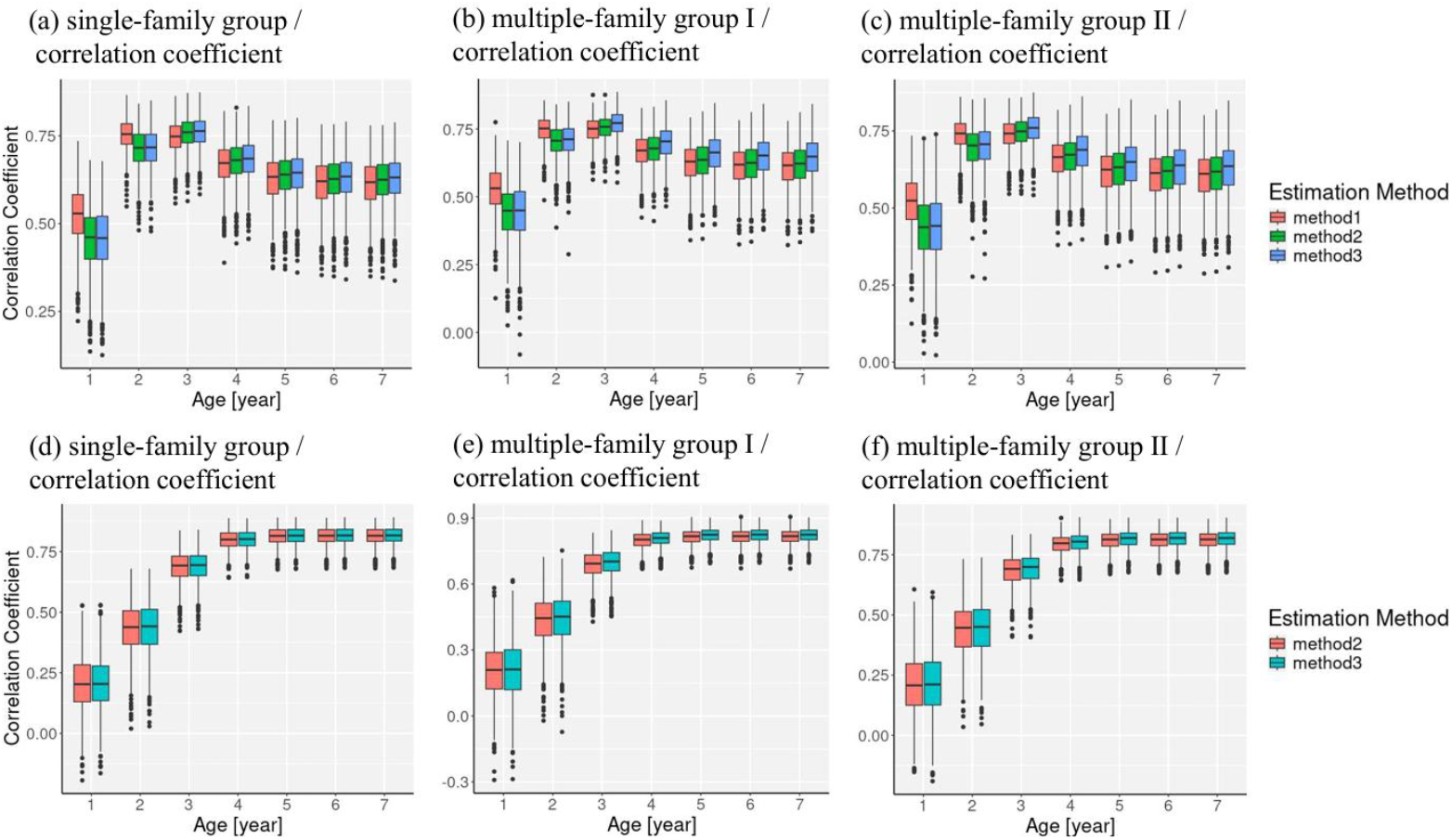
Estimation accuracy in each group. The upper three images (a), (b), and (c) show the results of Scenario 1, whereas the lower three images (d), (e), and (f) show the results of Scenario 2. Within each scenario, the left image corresponds to the single-family group, middle image to the multiple-family group I, and right image to the multiple-family group II. For each image, the estimation accuracy of the growth curve was evaluated annually using three methods, each represented by a distinct color. Box plots are not shown when the growth curve cannot be predicted.

## Discussion

This study explored the feasibility of estimating the logistic growth curve from incompletely fragmented longitudinal data using a Bayesian nonlinear model to impute missing values. This approach aimed to shorten the measurement period and reduce the burden of constructing a growth curve. The fragmented longitudinal data comprised three cohorts (youngest, middle, and oldest), and the growth curve was estimated using three different methods and groups across the two scenarios. Our focus is on the following three points: 1. Extent to which the estimation performance can be improved by considering data from other cohorts. 2. Whether the inclusion of genomic information enhances estimation performance. 3. How population structure impacts estimation performance. In addition, we evaluated the prediction performance of untested individuals based on the population structure.

### Improvements by leveraging data from individuals of other cohorts

A comparison of Methods 1 and 2 addressed the first question. While Method 1 requires estimating the growth curve using only data from a single cohort, Method 2 enables simultaneous estimation using data from all cohorts, leveraging information from individuals in other cohorts. The comparison was conducted in two scenarios, and we confirmed an improvement in the estimation performance for later growth in Scenario 1.

Method 2 in Scenario 1 was designed to enhance the estimation performance for later growth, particularly for parameter A, which characterizes later growth. Interestingly, the MSEs improved not only for A, but also for B and C. Although parameter A primarily represents later growth, its influence extends to the initial growth phase, where B and C play critical roles. Therefore, improving the estimation performance of parameter A reduces uncertainty in the initial growth phase, leading to better estimates of parameters B and C. This improvement is attributed to the properties of the nonlinear Bayesian model. The Bayesian nonlinear model assumes that the parameters of all individuals follow the same distribution, thus preventing the parameters from deviating excessively from the overall behavior. In Method 1, parameter A for the youngest cohort became highly unstable due to the absence of later growth data, whereas the middle and oldest cohort parameters were estimated more reliably. This assumption in the Bayesian nonlinear model helps bind all the parameters together, thereby stabilizing the estimation performance for the youngest cohort.

Such a function to impute missing values can be expected not only in the Bayesian nonlinear model but also in the NLMEM, where random effects are assumed to follow the same distribution. The application of the NLMEM to incomplete data with missing values has been documented in various studies^24,25^. Additionally, some studies such as ours have attempted to estimate later growth from initial growth data^32^. In a shift from plant-related research, a functional linear mixed model (FLMM) was employed in another study to estimate adult growth from children’s data, using previously obtained completed longitudinal data^33^. However, our study aims to shorten the measurement period, and is distinct in that it applies specifically structured missing data, known as fragmented longitudinal data.

Meanwhile, in Method 2 of Scenario 2, in which the growth curve of the older cohort was estimated using data from all cohorts, the estimation performance of parameters B and C, which characterize initial growth, dropped below 0.5. This decline can be attributed to the exclusion of the first-year data and uniform measurement intervals. We hypothesized that the targeted growth curve would represent the trunk circumference of the scion grafted onto the rootstock. Since measuring trunk circumference in the first year is difficult due to the thinness of the trunk, the first-year data were excluded from the fragmented longitudinal data, resulting in a lack of information on the initial growth phase. Additionally, assuming annual measurements over 6 years with constant intervals led to unbalanced sampling, as reflected in the distribution shift shown in Fig. 1B, where the initial growth was not sufficiently captured. The importance of the measurement interval has been well documented in the prior research^34^. Although our study indicates that estimating initial growth from fragmented longitudinal data is challenging, the performance can be improved by adjusting the measurement intervals.

### Improvement by genomic information

To explore the potential of genomic information for imputing missing values, Method 3 was designed by incorporating a genomic relationship matrix (GRM) into Method 2: Specifically, the improvement in the estimation performance by considering the GRM within the Bayesian nonlinear model was examined across three groups: the single-family group, multiple-family group I, and multiple-family group II. The superiority of Method 3 over Method 2 was confirmed, although the magnitude of superiority varied by group.

Generally, the longitudinal data at specific sampling points exhibit correlations with certain parameters. For example, late-stage longitudinal data are strongly correlated with parameter A, whereas initial-stage longitudinal data are more closely correlated with parameters B and C. Traditionally, parameter estimation has relied on these correlations. However, in this study, a large amount of data was missing, making it difficult to rely solely on correlation-based estimations. Nevertheless, when genomic information is considered, the parameters in the Bayesian nonlinear model are linked not only to phenotypes but also to genomic relationships; individuals who are genetically closer tend to have similar parameter values, and those who are genetically distant show more variation. In such cases, parameters can be estimated based on genomic relationships, which we believe will contribute to improvements in estimation performance. The algorithm for incorporating GRM into nonlinear models has been developed recently^29,30,35,36^, and to the best of our knowledge, this is the first study to demonstrate the potential of genomic information for imputing missing values within a nonlinear framework.

### Influence of population structure

While the inclusion of genetic information has been confirmed to improve the estimation of growth curves from fragmented longitudinal data, the magnitude of this improvement, as evaluated by accuracy, depends heavily on the group structure. Multiple-family group I showed a better improvement in estimation accuracy, whereas the single-family group yielded the poorest results. The key differences between these groups lie in the populations used to generate the longitudinal data and the strategy for allocating individuals to the three cohorts. Because both the single-family group and multiple-family group I used the same allocation strategies, their estimation performances can be compared based solely on the population structure. The genetic structure of the population used in this study, as visualized by PCA, clearly showed a wider range of genetic variation in the multiple-family group than in the single-family group. As we are evaluating the improvement guided by incorporating genomic information, a broader range of genetic variation suggests great potential for improvement, which likely contributes to the higher accuracy in multiple-family group I.

Meanwhile, multiple-family Groups I and II used the same population to generate longitudinal data but applied different allocation strategies, allowing for the comparison of estimation performance based solely on allocation patterns. Although the allocation strategy used in multiple-family group II was more realistic, its estimation performance was lower than that of multiple-family group I. When individuals are randomly assigned, as in the multiple-family group I, genetically related individuals, such as those from the same family, are distributed more evenly across the three cohorts, making it easier to estimate the growth curve. In contrast, when individuals from the same family are concentrated in a specific cohort, such as in the multiple-family group II, the phenotype data across cohorts seem less related, making it more difficult to estimate phenotypic values across cohorts. In genomic selection, the prediction performance is strongly influenced by the genomic relationship between the training and test populations, leading to numerous studies on the impact of the allocation strategy^37^. However, the experimental setup described in this section differs in that it focuses on longitudinal genetic relationships rather than on the relationship between the training and test populations. Another key distinction is that this section emphasizes estimation rather than prediction. The prediction performance considering the allocation of the training and test populations is discussed in the subsequent section.

Prediction of untested individuals

Although our primary focus has been on the estimation performance of tested individuals, the incorporation of the GRM into the Bayesian nonlinear model also enables the direct prediction of untested individuals. In this study, we attempted to build a model using fragmented longitudinal data and evaluated the prediction performance for each group. However, the prediction performance was low, particularly for parameters B and C; even parameter A did not exceed 0.4, indicating the challenge of prediction from fragmented longitudinal data.

Among the three groups, the single-family group and multiple-family group I exhibited the best prediction performance for parameter A, whereas multiple-family group II exhibited the poorest performance across all parameters. The relationship between the prediction performance and population structure has been actively studied in the context of genomic selection. For example, a previous study categorized a wheat population based on training and test population structures and found that prediction performance declined when the family of the test population was excluded from the training population, attributing the decline to the localization of family specific genes^37^. Although the prediction performances of B and C were insufficient, they may be improved by optimizing the sampling time, as mentioned earlier, which warrants further investigation.

## Materials and Methods

### Plant material

In this study, we used 505 individuals with different genotypes from 11 families generated by the National Agriculture and Food Research Organization, Institute of Fruit Tree and Tea Science (Okitsu, Shizuoka, Japan). The parents of these families had diverse backgrounds, including tangerines, lemons, and pomelos, and the families were numbered from 1 to 11. Family 11 comprised 266 progeny, whereas the remaining 10 families had an average of 20 individuals.

### SNP genotyping data

We conducted GRAS-Di^38^ analysis using Illumina HiSeq 4000, generating 150 bp paired-end reads according to the manufacturer’s protocol, with an average of approximately 8 million reads per sample. We then mapped the trimmed clean reads to the clementine haploid reference genome v1.0 (https://data.jgi.doe.gov/refinedownload/phytozome?organism=Cclementina&expanded=Phytozome-182) using GSNAP (version 2021-12-17)^39^ with options -k = 15 and-max-mismatches=0.125, resulting in BAM files for individual samples. After sorting the BAM files in coordinate order using Picard tools, the secondary mapped reads were marked and index files were generated. We called polymorphic sites using the GATK4 (4.5.5.0) HaplotypeCaller and created GVCF files for each sample. Next, we merged the GVCF files using GATK4 CombineGVCFs, and converted the merged GVCF file into a VCF file using GATK4 GenotypeGVCFs. The VCF file was filtered with thresholds of meanDP >= 20 and meanGQ >= 20 using vcftools^40^. Imputation was performed using Beagle 5.2^41^, selecting data based on a minor allele frequency (MAF) > 1%. Using Picard, we verified the Mendelian violations in 387 trios and excluded loci with inconsistencies in more than two trios. Finally, the linkage disequilibrium was calculated, and SNPs were selected based on a linkage disequilibrium threshold of less than 0.95 using “LD.thin” function of the R package RAINBOWR^42^, resulting in a total of 45,929 makers.

### Population structure analysis

Principal component analysis was performed using the SNP genotyping data of 505 individuals with the “prcomp” function of the R package stats.

### Longitudinal data generation

The longitudinal data were generated using a growth model. Among various growth models, such as the Gomperz, von Bertalanffy, and Richards models, the logistic model was best suited to obtain longitudinal data of citrus trunk circumference of citrus breeding populations 2 to 7 years after grafting onto 3-year-old trifoliate orange rootstock (data not shown); therefore, it was used in our study. The logistic model was parameterized as follows:

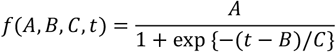

where, *A* is the asymptotic parameter, represents the maximum value of the growth curve, B is the inflection time, and *1*/*C* is the growth rate at the inflection point. We assumed that these parameters were unique to each individual, and the unique parameters *A*_*i*_, *B*_*i*_, and *C*_*i*_ for the *i*-th individual were determined using the following equations:

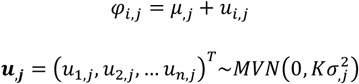

where φ_*i,j*_ is the *j*-th parameters of *i*-th individual, *μ*_,*j*_ is the mean of the *j*-th parameters, and *u*_*i,j*_ is the deviation from the global mean, characterizing each individual. 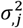 is the variance and *k* is the genomic relationship matrix, calculated for both the single-family group and the multiple-family group using the “calcGRM” function of the RAINBOWR package. In the process of data generation, the mean *μ*_,*j*_ and the variance 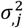 were first fixed, and then the individual-specific parameter φ_*i,j*_ was randomly generated. These parameters were applied to a logistic model to determine individual-specific growth curves. The values of *μ*_,*j*_ and 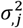 were determined to mimic the actually measured citrus trunk circumference growth data as follows:

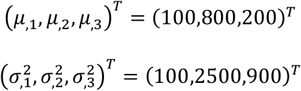

Data for each year from 2 to 7 years were extracted from the generated individual-specific growth curves, and longitudinal data were completed by adding measurement noise. To ensure that heritability (i.e., the proportion of variance explained by genetic factors) remained constant over time, the variance of the longitudinal data was first calculated at each sampling point, and then the magnitude of variance was adjusted to maintain a heritability of 0.5.

### Fragmented longitudinal data generation

Fragmented longitudinal data were obtained from the artificially generated longitudinal data. Specifically, the 4th to 7th years’ data were replaced with missing values for the youngest cohort, while the 2nd, 3rd and 6th, and 7th years’ data were replaced with missing values for the middle cohort. Similarly, the 2nd to 7th years’ data were replaced with missing values for the oldest cohort.

### Estimation of latent variable

To estimate the three parameters, *A*_*i*_, *B*_*i*_, and *C*_*i*_, of the logistic model, we applied a Bayesian nonlinear model. Individual growth curves were estimated by incorporating these parameters into the model functions. The Bayesian nonlinear model is defined as follows:

Let *f* represent a nonlinear growth model, ***D*** the model input, ***ϕ*** the model parameters, ***Y*** the observations, and *i* the genotype index. The model can be expressed as follows:

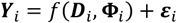

where *ε*_*i*_ is the error term, assumed to follow a multiple normal distribution:

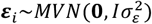

The prior distribution of the residual variances 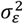 is the Jeffreys’ scale-invariant prior:

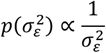

The *j*-th parameter of *ϕ*_*i*_ is regressed on genome-wide marker genotypes as:

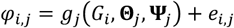

where *G*_*i*_ is the marker genotypes, *g*_*j*_ is the whole-genome regression function, Θ_*j*_ is the regression parameters of *f*_*j*_, Ψ_*j*_ is the hyperparameters of *f*_*j*_, and:

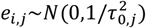

The prior for 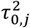 follows

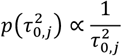

In this study, we used a GBLUP model for whole-genome regression, defined as

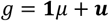

with priors:

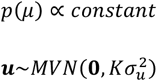

where *k* is the additive genomic relationship matrix. The prior for *σ*^2^ is:

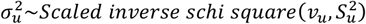

The parameters to estimate Θ are 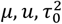, and 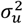, and the hyperparameters Ψ are *ν* and 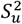. For Methods 1 and 2, in which genomic information was not used, the additive genomic relationship matrix *k* was replaced by the identity matrix *I*.

Estimation was performed using the “GenomeBasedModel” function of the GenomeBasedModel package in R, with the initial value set by the “drm” function of the drc package^43^. In the initial value calculation, the longitudinal dataset of all individuals was treated as a single individually derived dataset. Iterations were run 1000 times for each generated dataset, and the estimation was evaluated using both Pearson’s correlation coefficient and the mean square error.

### Prediction of untested individuals

When genomic information is incorporated into the Bayesian nonlinear model, the estimated parameters follow a multivariate normal distribution with a variance-covariance matrix proportional to the GRM. In this case, the logistic model parameters of untested (predicted) individuals, 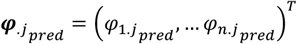, can be predicted based on the genomic relationship matrix and the estimated parameters of tested (observed) individuals, 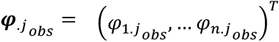, using the following equation^44^:

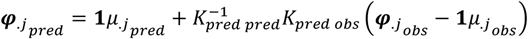

where *k*_*pred pred*_ is the additive genetic relationship matrix corresponding to the predicted individuals, and *k*_*pred obs*_ is that of the predicted and observed individuals. 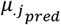 and 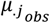 are the means of the predicted and estimated parameters (i.e., estimated parameters of an observed individual) respectively. In predicting 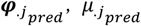 is replaced by 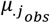. A growth curve was then constructed using these parameters.

To evaluate the prediction performance of the model constructed using the fragmented longitudinal data, we conducted the following experiment: For the single-family and multiple-family group I, one eleventh of the individuals were used as the training population, and the rest were used as the training population, which was further divided into three cohorts. Cross-validation was performed by rotating the test population 11 times and the prediction performance was calculated. This process was repeated 1000 times using different longitudinal data. In the case of multiple-family group II, individuals were allocated by family, and each family was selected as the test population for each rotation. It should be noted that because all individuals across the three cohorts were used for growth curve prediction, there was no distinction between Methods 1 and 2.

### Significance test of the difference in prediction performance

Because the variance in prediction performance across all simulation iterations differed by group, a statistical significance test for prediction performance was conducted using the Steel-Dwass asymptotic test. This was implemented with the “pSDCFlig” function in “NSM3” package in R.

## Acknowledgments

We are grateful to all members of the National Agriculture and Food Research Organization Institute of Fruit Tree and Tea Science for maintaining the Citrus trees, as well as Kosuke Hamazaki and Kengo Sakurai for sharing their expertise in the simulation analysis. This research is supported by a grant from MAFF commissioned project study on “Smart breeding technologies to Accelerate the development of new varieties toward achieving “Strategy for Sustainable Food Systems, MIDORI”” and Japan Science and Technology Agency – OPERA (Program on Open Innovation Platform with Enterprises, Research Institute and Academia) Grant Number JPMJOP1851.

## Data Availability Statements

Data available on request.

## Conflict of interests

The authors declare that they have no conflict of interest.

## Competing financial interests

The authors declare no competing financial interests.

## Author Contributions

S. K. and H. I. conceived and designed the study. K.N. designed the study. T.S. extracted DNA and performed SNP genotyping. S.K., M.F.M., T.S., H.I., and K.N. conducted phenotyping. S. K. performed the simulations. H.I. provided technical help for statistical analysis. S. K. and T. S. drafted the manuscript. All the authors have read and approved the manuscript.

## Notes

### Competing Interest Statement

The authors have declared no competing interest.

